# Combining AI structure prediction and integrative modelling for nanobody-antigen complexes

**DOI:** 10.1101/2025.07.01.662355

**Authors:** Miguel Sánchez-Marín, Marco Giulini, Alexandre M.J.J. Bonvin

## Abstract

Nanobodies exhibit antigen binding affinities of the same order as those of antibodies, which, along with their small size and unique structural characteristics, makes them well-suited for therapeutic and diagnostic applications. The lack of coevolutionary signals in nanobody-antigen complexes together with the broad complementary determining region 3 loop (CDR3) conformational space poses a challenge for predicting the 3D structure of those complexes with computational modelling and artificial intelligence-based methods. In this context, physics-based information-driven docking can provide an alternative solution. This study evaluates the state-of-the-art of machine learning-based methods for nanobody structure prediction and benchmarks various HADDOCK workflows to model their interaction with antigens using different input nanobody ensembles and information scenarios. We propose an ensemble docking pipeline that achieves high success rates starting from nanobody structural models predicted by AlphaFold2 and ImmuneBuilder. Our results highlight the effectiveness of physics-based complex prediction of immune proteins when accurate input structures and sufficient information to guide the modelling are available.

## 1 Introduction

Heavy-chain antibodies are a special type of antibodies that lack the light chain and the first constant region of the heavy chain. This kind of antibodies are present in the serum of camelids in addition to canonical antibodies [1]. Single-domain antibodies, or nanobodies, consist of the single variable domain from the heavy chain corresponding to the smallest possible independent antigen binding unit of heavy-chain antibodies [2]. The exceptional structure of nanobodies entails interesting features such as high solubility, high stability, and a binding affinity comparable to that of conventional antibodies, thus making them suitable candidates for therapeutic and diagnostic applications [3, 4, 5, 6, 7].

The nanobody structure resembles that of canonical antibodies VH regions, with four conserved Framework Regions (FR1-FR4) and three intercalated hypervariable loops forming the Complementarity Determining Regions (CDR1-CDR3). CDR loops are responsible for antigen binding and are optimized for this function during maturation by mutation processes [2, 8, 9]. In nanobodies, CDR3 is usually longer and more flexible than in antibodies, resulting in a very diverse conformational space at the structural level [10].

Two broad classes of CDR3 conformations have been described: ‘kinked’, where the loop folds back and interacts with the framework regions to counteract the absence of the light chain (the most frequent ones), and ‘extended’, where this folding does not occur and the framework regions are more exposed [11, 12]. Nanobodies exhibit additional differences from VH structures, such as substitutions in FR2 residues increasing hydrophilicity, and often a non-canonical disulfide bond between CDR3 and CDR1 or CDR2. These differences are thought to account for the lack of a light chain to stabilize the structure [13, 14, 15].

These structural differences result in a slightly different binding mode to antigens between nanobodies and antibodies, with CDR3 usually considered the main contributor to binding in nanobodies: its increased length and variability allow for epitope shape complementarity and large paratope contributions [10, 16]. In addition, the nanobody framework regions contribute more to the binding because of the absence of a light chain. The interacting framework residues are conserved and belong to all framework regions, with FR2 being the most frequent framework interactor [10, 16, 17].

Understanding nanobody-antigen interactions at a structural, atomistic level is of utmost importance given their potential therapeutic applications. Information about their binding mode can be extracted from a variety of experimental approaches such as for example mutagenesis, hydrogen/deuterium exchange, nuclear magnetic resonance, or X-ray crystallography [18, 19, 20, 21]. Experimental approaches are, however, usually expensive and time-consuming, impeding high-throughput research.

In recent years, computational approaches have arisen as powerful alternatives. Several machine learning-based approaches have been developed for accurate 3D structure prediction of immune proteins. AlphaFold2 [22], AlphaFold2-Multimer [23], and related approaches [24, 25] typically use coevolutionary signals encoded in multiple sequence alignments for the prediction of protein and protein-protein complexes. Their high performance usually comes at rather high computational costs, and the prediction of orphan proteins and non-coevolved complexes remains challenging for these methods [26]. Deep-learning methods specifically trained on nanobody data such as ImmuneBuilder [27] and NanoNet [28], together with language model-based methods such as RaptorX-Single [29], try to address these limitations by not relying on MSAs, reducing the computational cost associated with nanobody structure prediction. Even so, modelling the nanobody CDR3 loop remains challenging given its high sequence variability and inherent flexibility [10, 30].

Similarly, the prediction of nanobody-antigen complex faces challenges due to the absence of coevolutionary signals, limiting the accuracy of deep-learning protein-protein complex prediction methods for this kind of complexes [26], even if some immune-specific algorithms have been developed with this purpose [31]. AlphaFold3 [32] reports a large improvement in antibody-antigen complex prediction success rates (SR), reaching up to 60% acceptable quality models on antibody-antigen complexes, but this at the cost of an increased sampling strategy (1,000 seeds / 5,000 models). Despite this improvement, the required number of sampled models is computationally very demanding and therefore not broadly usable. Most of these methods provide confidence metrics that are highly correlated with the quality of the prediction, thus enabling users to easily determine when they can be trusted.

Given its proved success in modelling antibody-antigen complexes [33, 34], information-driven, physics-based docking with HADDOCK [35] appears as a reasonable solution to tackle the nanobody-antigen complex prediction challenge in cases where some information is available to drive the process. With the aim of designing a reliable nanobody-antigen prediction pipeline, we first assembled a non-redundant dataset of nanobody-antigen crystal structures, avoiding any data leakage with the training sets used by the various AI prediction methods. We then assessed the performance of state-of-the-art, AI-based monomeric and complex immune protein prediction algorithms. We analyse the quality of the predicted (unbound) structures and use those as input for benchmarking different docking protocols in HADDOCK considering various levels of information.

We demonstrate how HADDOCK can provide acceptable nanobody-antigen models coming from AI-predicted monomers. Our modelling pipeline combines nanobody models generated with AlphaFold2-Multimer and ImmuneBuilder as input for HADDOCK. To harvest the information about the binding mode of nanobodies, we propose two alternative paratope representations to guide the docking process, which also involve framework information, to guide the docking process. This results in a substantial improvement in the performance of the docking. The pipeline is robust across various levels of paratope and epitope information, making it highly promising for nanobody-focused therapeutic research.

## 2 Results

We have defined a benchmark dataset consisting of 40 nanobody-antigen complexes, non-redundant in their CDR3 sequences (see Supplementary Table 1), released after the cutoff date of the used methods (September 30th, 2021, see Methods) and without any sequence homology in the CDR3 region with other nanobodies seen by the various AI predictors. The targeted antigens come from viruses (17 antigens), human (8), bacteria (7), rodents (4), plant (3) and protozoa (1) (see Supplementary Table 2).

### 2.1 AlphaFold has moderate accuracy in nanobody-antigen complex prediction

We first assessed the performance of AlphaFold2-Multimer v.2.3 (AF2M) on the benchmark dataset. To evaluate the quality of the complex predictions we compare the acceptable/medium/high success rate (SR) for the topN ranked models, using the CAPRI quality assessment criteria [36, 37] (see Methods) and also report DockQ [38], a continuous metric that combines the CAPRI criteria into a score ranging from 0 to 1.

AF2M achieved a mean DockQ of 0.211 for first-ranked models, with an acceptable quality SR on Top1 predictions of 25.0% (15.0% medium and 10.0% high). When considering the Top10 predictions, the SR only slightly increased to 27.5% acceptable, of which 17.5% medium and 15.0% high. Our results do confirm the limited performance of AF2M in nanobody-antigen complex predictions in comparison with other protein-protein complexes, consistent with previous studies [26, 31].

During the time course of this work the AlphaFold3 server (AF3) was released, reporting an impressive improvement in nanobody-antigen complex predictions [32]. On our dataset, AF3 (25 models generated per complex) has a mean DockQ of Top1 models equal to 0.324, with an acceptable quality SR on Top1 predictions of 32.5% acceptable (32.5% medium and 25% high), and reaches 52.5% when considering the Top10 predictions (47.5% medium and 30% high). The limited number of seeds used results in lower SR from that reported in Abramson *et al*. [32] (60 % Top1 SR, obtained with an extensive sampling strategy with 1000 seeds - 5000 models), consistent with other benchmarking studies [39, 40]. AF3 certainly represents a major leap in structural accuracy for *ab-initio* nanobody-antigen complex modelling.

### 2.2 Combining different prediction methods improves the accuracy of the nanobody models

We assessed the accuracy of six different methods for the structure prediction of nanobodies, namely AlphaFold2.3-Multimer (AF2M), Alphafold2.3 (AF Monomer, AF2 from now on), ImmuneBuilder (IB), NanoNet (NN), RaptorX-Single (RXS), and Alphafold3 (AF3). All methods predict the framework region with a mean RMSD below 2Å and CDR1/CDR2 below 2.5Å (see Table 1). Unsurprisingly, CDR3 proved to be the most challenging region to predict, with all methods showing mean RMSDs above 3Å. In the following, we will use the CDR3 RMSD as the quality evaluation metric. All RMSDs were calculated on the backbone atoms of the defined regions (e.g. CDR3), after fitting the predicted structure onto the reference on the framework regions.

**Table 1:** Achieved backbone mean RMSD (± standard deviation) of the different nanobody regions by the different prediction methods. The machine learning based nanobody structure prediction methods AlphaFold2-Multimer (AF Multimer), AlphaFold2 (AF Monomer), ImmuneBuilder (IB), RaptorX-Single (RXS), NanoNet (NN) and AlphaFold3 (AF3) performances are shown. For methods generating several models, only the first ranked model was considered.

| Prediction<br>Method | Framework<br>RMSD (Å) | CDR1<br>RMSD (Å) | CDR2<br>RMSD (Å) | CDR3<br>RMSD (Å) |
| --- | --- | --- | --- | --- |
| AF Multimer (top 1) | 1.05±0.66 | 2.14±1.45 | 1.95±1.24 | 3.10±1.97 |
| AF Monomer (top 1) | 1.04±0.63 | 1.94±1.22 | 1.83±1.07 | 3.39±2.11 |
| IB (top 1) | 0.86±0.39 | 1.89±1.21 | 1.83±1.07 | 3.23±1.82 |
| RXS | 1.04±0.70 | 2.13±1.50 | 2.00±1.40 | 3.28±2.27 |
| NN | 1.89±1.53 | 2.48±1.42 | 2.35±1.30 | 4.60±2.63 |
| AF3 (top 1) | 1.03±0.69 | 1.92±1.71 | 1.76±1.35 | 2.48±2.32 |

The best performing method in terms of CDR3 RMSD of the first ranked models was AF3, with a mean CDR3 RMSD of 2.48Å (see Table 1). AF2M, IB, RXS, and AFMo follow with mean RMSDs of 3.10Å, 3.23Å, 3.28Å, and 3.39Å, respectively. NN performs worse than the other predictors, with a mean CDR3 RMSD of 4.60Å. With the exception of AF3, the differences between the best methods are not significant considering the large standard deviations (see Table 1). All distributions are shown in Supplementary Fig. 1. The performance of AF2M, IB, and NN is comparable to previously reported results [31, 41, 42].

Since the conformational space of the nanobody loops can potentially be very large, a broader sampling of models can be highly useful for any downstream modelling task, including docking. We therefore evaluated the best prediction from all generated models per method to measure the maximum achievable nanobody accuracy. AF3 outperforms all the other methods with a mean and median CDR3 RMSD of 1.84Å and 1.15Å, respectively (Fig. 2a). As the restricting licensing conditions do not allow us to perform any downstream modelling with AF3 structures (from the AF3-server), we did not consider them for further analysis in this work.

**Figure 1:**
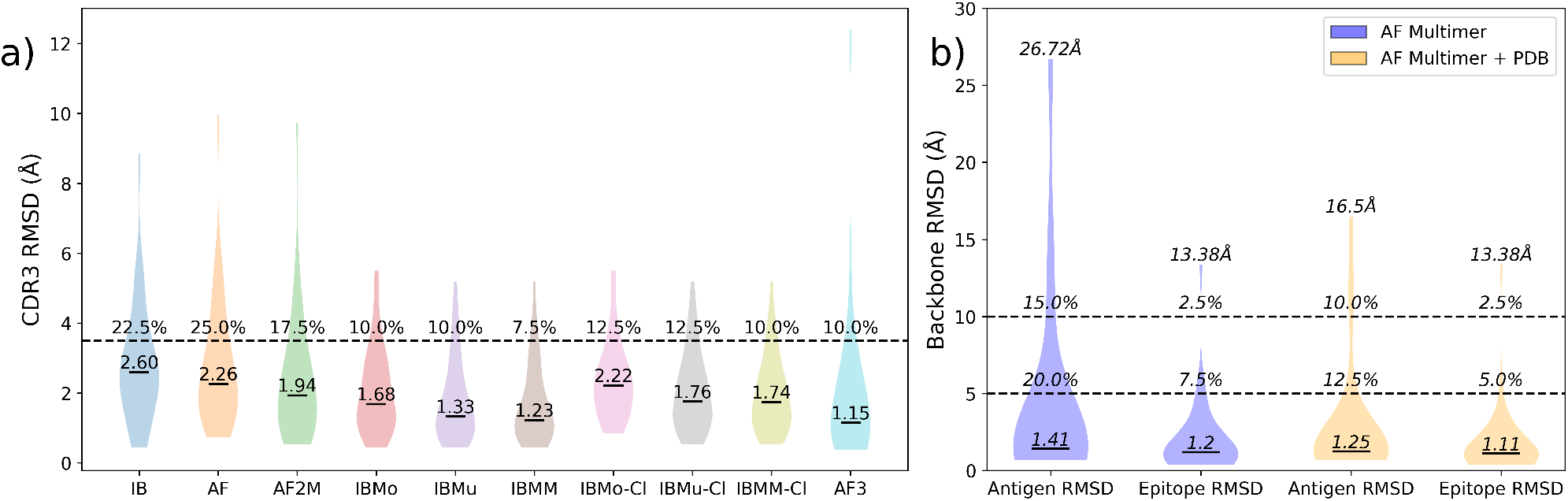
Accuracy of the predicted unbound nanobody and antigen structure ensembles. **a)** Violin plot showing the distribution of the best backbone CDR3 RMSD in the ensembles coming from multiple prediction methods ImmuneBuilder (IB), AlphaFold2 (AF2), AlphaFold2-Multimer (AF2M) and AlphaFold3 (AF3); the combination of these ensembles (IBMo=IB+AF2, IBMu=IB+AF2M, IBMM=IB+AF2+AF2M); and the combined ensembles after clustering (IBMo-Cl, IBMu-Cl, IBMM-Cl). Medians are shown in the plot. The dashed line and the numbers above show the percentage of dataset entries for which no structure in each ensemble have CDR3 RMSD ≤ 3.5Å **b)** Distributions of the full antigen and epitope backbone RMSD for the best AlphaFold2-Multimer predicted unbound antigen dataset (blue) and for the dataset including PDB homologous structures (orange, see main text). Medians are shown in the plot, together with the highest antigen and epitope RMSD values (on top). The percentages of structures with antigen or epitope RMSD higher than 5 and 10Å are reported above the dashed lines.

**Figure 2:**
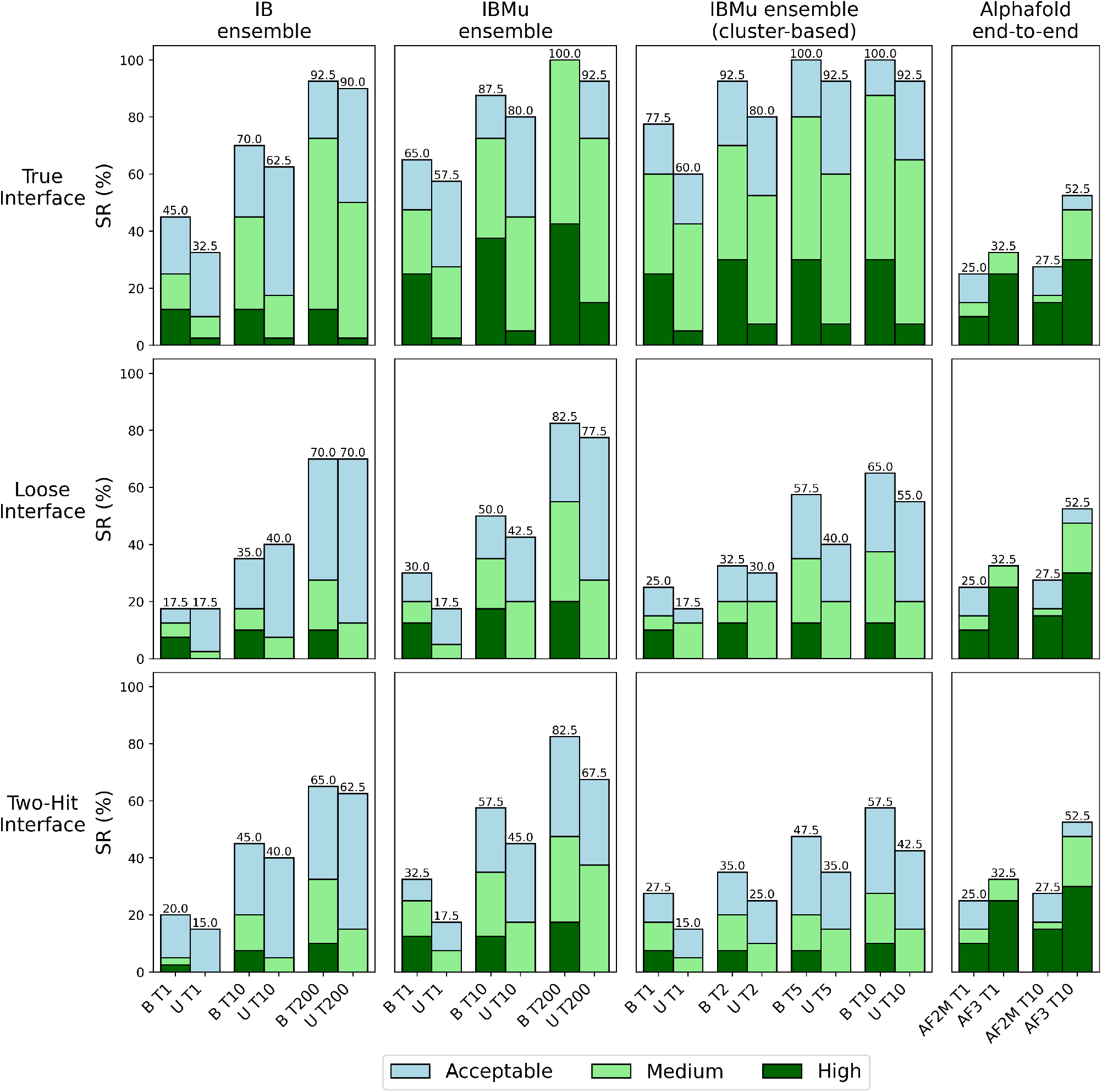
HADDOCK3 performance on nanobody-antigen docking. HADDOCK success rates (SR) for nanobody-antigen docking models after Flexible Refinement stage for True, Loose, and Two-Hit Interface scenarios. Docking success is reported for the ensembles ImmuneBuilder (IB) and ImmuneBuilder + AlphaFold2-Multimer (IBMu), both for bound (B) and unbound antigens (U) and calculated for Top 1 (T1), Top 10 (T10) and Top 200 (T200) ranked models. For IBMu we report also the cluster-based success rate over T1, T2, T5 and T10 clusters. Clusters were defined based on Fraction of Common Contact (FCC) clustering [46] of the models at the end of the workflow. Success is considered when at least one of the four first-ranked models in any of the TopN ranked clusters is classified as acceptable/medium/high quality. The AlphaFold2-Multimer (AF2M) and AlphaFold3 (AF3) SR are provided on the right. Light blue, light green, and dark green indicate acceptable, medium, and high-quality models, respectively, as defined by the CAPRI criteria. Acceptable SR (%) are shown on top of the bars.

The AF2M ensemble (25 models per case) has the second-best accuracy with a median CDR3 RMSD 1.94Å on the benchmark dataset (see Fig. 2a), followed by the IB ensemble (4 models) and AF2 ensemble (25 models). The combination of these ensembles substantially improves the accuracy of the best achievable prediction, with the best combination corresponding to the ensemble merging all the IB, AF2, and AF2M models with a median CDR3 RMSD of 1.23Å (Fig. 2a).

As the large number of models in the combined ensembles could lower the docking success, we performed hierarchical clustering based on the CDR3 RMSD and selected the best model from each cluster based on confidence metrics (see Methods). After clustering, the clustered ensembles combining different methods are still more accurate than the single-method ones. The best combination is obtained from clustered IB and AF2M model ensembles (median CDR3 RMSD = 1.76Å). Adding AF2 models doesn’t lead to a significant improvement (median CDR3 RMSD = 1.74Å, Fig. 2a).

In a recent work [43], we benchmarked the effectiveness of AlphaFlow (AFL) [44] to increase the diversity and accuracy of H3 loops of antibodies, demonstrating how clustering AFL models can be useful to obtain loop conformations with lower CDR3 RMSD with respect to the experimental structure. Here we follow the same approach, sampling 1000 AFL models for the nine nanobodies for which neither AF2 nor AF2M provided any model with CDR3 RMSD ≤ 3Å. Following Ref. [43] we clustered AFL models in a clustered ensemble of 20 clusters and checked the accuracy of the best model within this ensemble. In contrast to antibodies, AFL does not seem to provide any advantage in this case. The clustered ensemble only provides substantially better loop conformations (improvement in CDR3 RMSD ≥·1.0Å) in two cases, one of the two being PDB ID 7TPR, for which the best AFL conformation is still utterly incorrect (CDR3 RMSD = 7.20Å), although better than the AF2 and AF2M models.

### 2.3 The quality of antigen structure prediction with AlphaFold2-Multimer can be limiting for docking

We selected the best antigen structures from the AF2M complex predictions, based on the global predicted local distance difference test (pLDDT). Quality assessment showed that 9 structures presented a global RMSD above 5Å and 3 structures an epitope RMSD above 5Å (Fig. 2b). As poor accuracy antigen structures are a known limiting factor for docking [34], we performed a sequence search in the PDB in order to find homologous antigen structures when available (see Methods). For 24 of the unbound antigens, homologous structures could be found in the PDB. For the remaining 16 antigens, the best AF2M predictions were selected (see Supplementary Table 2). The resulting combined dataset has only 5 structures showing a global RMSD above 5Å and 2 with an epitope RMSD above 5Å.

In cases where the epitope spanned multiple antigen chains, the relative orientation between chains proved to be the limiting factor in AF2M, as previously noted [45]. For one of these multiple antigen chain cases (PDB ID 7QNE), no similar structures were available in the PDB, hindering the generation of reasonable starting unbound models (global and epitope RMSD *>* 10Å). This led us to consider this case as failed for all unbound docking experiments.

### 2.4 Information-driven HADDOCK unbound docking

We assessed the docking performance of HADDOCK starting from different unbound nanobody ensembles targeting the modelled antigens. HADDOCK allows the incorporation of restraints based on binding site information of the components of a complex to guide the docking process. With the aim of evaluating HADDOCK’s performance, we defined four different scenarios based on the possible information available: True Interface scenario (TI, ideal scenario where all interface residues are known, but not the contacts they are forming), Loose Interface scenario (LI, only a loose definition of the epitope is available and no information on the nanobody, except for the knowledge of the CDRs), Two-Hit Interface scenario (2I, only two epitope residues, key for the stability of the nanobody-antigen complex as identified, for example, by mutagenesis, are available) and All Surface scenario (AS, no information on the interface of the antigen is available).

We applied the default HADDOCK antibody-antigen docking pipeline [33], with some sampling adaptations for the AS scenario (see Methods). We analyzed four different ensembles of unbound nanobody structures: the combined methods ensembles (IBMu, IBMo, IBMM) after clustering, and the ImmuneBuilder ensemble as a “computationally fast” option. We tested the unbound nanobody ensembles against both bound and unbound (predicted) antigen structures, with the objective of analyzing the impact of predicted antigen models on the docking performance.

Among all scenarios, the best performances were almost always achieved at the Flexible Refinement stage of the pipeline, so we focused on the results obtained at this stage. TI runs achieved high SR, with up to 65% acceptable SR for Top1 and 87.5% for Top10 models in bound antigen runs, and 57.5% for Top1 and 80% for Top10 in unbound antigen runs (Fig. 3a). LI runs reached a maximum acceptable SR of 30% for Top1 models and 55% for Top10 in bound antigen runs, and up to 17.5% for Top1 and 42.5% for Top10 in unbound antigen runs. The 2I scenario showed very similar SR.

**Figure 3:**
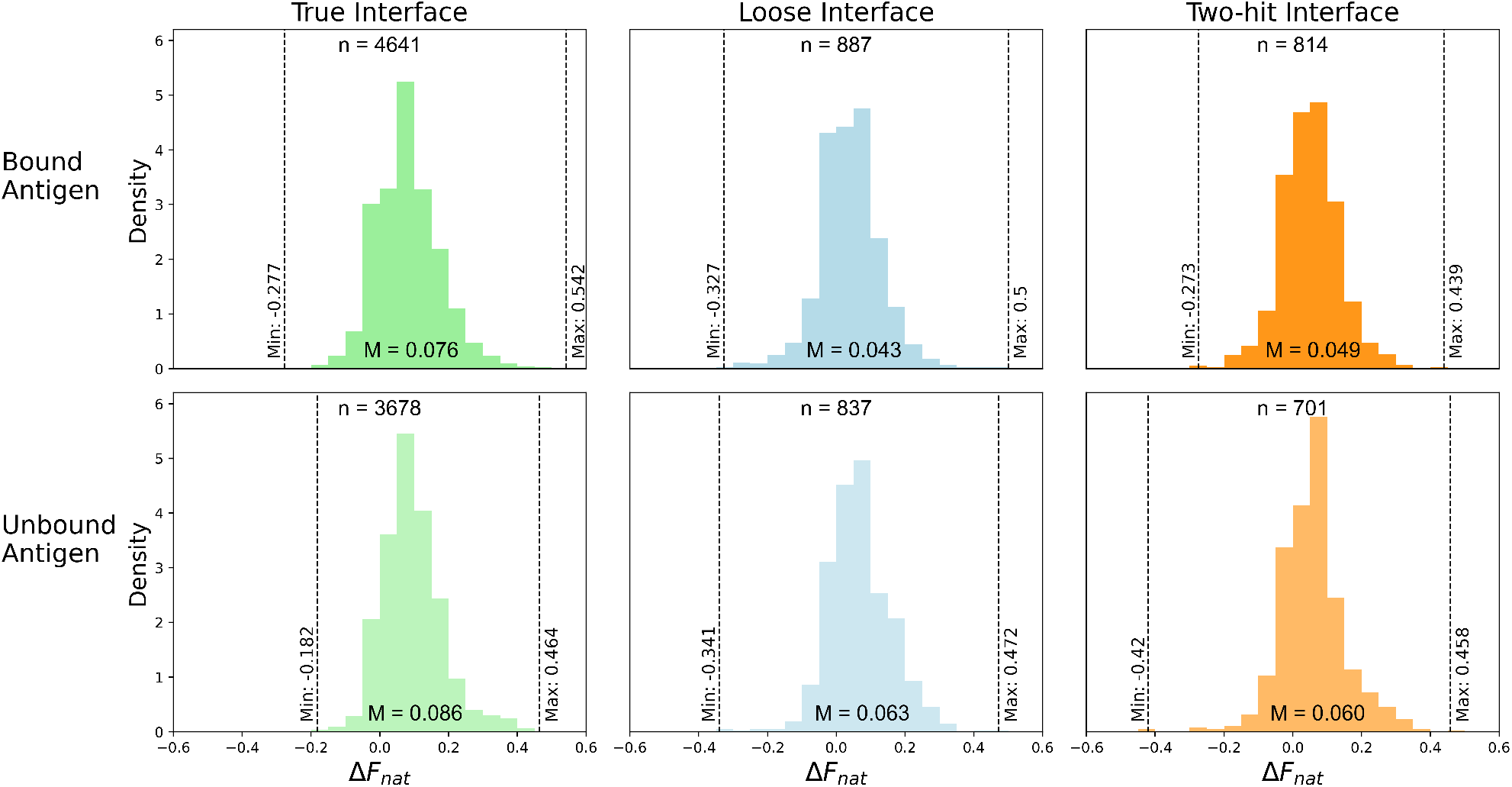
Fraction of Native Contacts improves after Flexible Refinement. The histograms show the distribution of changes in Fnat after flexible refinement 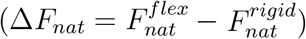 for all acceptable models (positive values indicate an improvement in the recovery of native contacts). Frequencies are shown for True, Loose and Two-Hit interface scenarios runs with bound and unbound antigen. The median Δ*F*_*nat*_ value is shown at the bottom of the histogram and maximum and minimum Δ*F*_*nat*_ are reported with black dashed lines. The number of models in the plotted distributions (n) is shown on each graph.

When looking at medium-quality models, bound antigen runs achieved 72.5%, 37.5%, and 35% SR, respectively. Medium SR are lower for unbound runs, with a Top10 SR of 45% for the TI scenario, while LI and 2I generate medium-quality models in Top10 only for 15-20% of the cases. Due to its high computational cost, we ran the AS scenario only on the best performing ensemble (IBMu), obtaining a modest 7.5% acceptable SR for Top1 and 20% for Top10 models in bound antigen runs (see Supplementary Fig. 2). Considering this low performance, AS scenario was not run for unbound antigen.

The use of unbound, predicted antigen models decreased the docking success with respect to the bound form along all ensembles and scenarios. The quality of the antigen modelling is thus critical for the success of nanobody-antigen docking. Maximum SR were frequently observed for the IBMu ensemble runs (Fig. 3a). This ensemble led to the best nanobody predictions among all ensembles (Fig. 2a). HADDOCK’s ability to generate good models is thus improved when more accurate structures are available in the starting ensemble, boosting the docking success.

The True Interface scenario achieved very high SR for all ensembles, demonstrating HADDOCK’s ability to generate good models when extensive information is provided. The Loose and Two-Hit Interface scenarios have lower SR, but still higher than AlphaFold2-Multimer (both for acceptable and medium SR, see Fig. 3a), demonstrating that, even when limited information on the epitope is available, this ensemble docking approach achieves better results than AlphaFold2-Multimer and comparable to those of AlphaFold3.

To further analyse HADDOCK’s performance compared to AlphaFold, we calculated its acceptable SR obtained for cases where AlphaFold2-multimer and AlphaFold3 did not provide acceptable models within their top ten predictions (see Table 2). Additionally, adopting a realistic approach in which the reference structure was not known, we used AlphaFold’s interface predicted TM score (ipTM [23, 32]) as a proxy for its success. Following this approach, we extracted the structures for which the highest AlphaFold ipTM is lower than 0.6 and measured HADDOCK success over those instances. Table 2 shows how HADDOCK can be a helpful resource in this context, providing an accurate solution in roughly 30/40% of the cases in the low information scenarios (LI and 2I). In the TI scenario, HADDOCK succeeds in the majority of cases, with failures typically occurring only when the input nanobody or antigen were severely mispredicted.

**Table 2:** HADDOCK acceptable success rate (acc. SR) on cases that are difficult for AlphaFold. Instances for which AlphaFold2-multimer and AlphaFold3 lack an acceptable model (according to CAPRI criteria) in the first 10 ranked models are displayed (AF2 Failed, AF3 Failed). Additionally, nanobody antigen complexes where AlphaFold’s maximum ipTM *<* 0.6 are shown (AF2 Low ipTM, AF3 Low ipTM). HADDOCK acceptable SR are reported across the three scenarios, as calculated over the best ranked model of the IBMu ensemble run (see main text). Statistics calculated for the first ten models are provided between square brackets.

| Dataset | # PDB | 2I acc. SR | LI acc. SR | TI acc. SR |
| --- | --- | --- | --- | --- |
| AF2 Failed | 29 | 7 [38] | 10 [34] | 52 [72] |
| AF2 Low ipTM | 27 | 15 [48] | 19 [41] | 63 [85] |
| AF3 Failed | 18 | 17 [28] | 22 [39] | 61 [83] |
| AF3 Low ipTM | 21 | 14 [29] | 29 [38] | 52 [81] |

We observed high acceptable SR when looking at all models (Top200) in comparison with Top1 and Top10, stressing the difficulty to properly score and identify the best models in this case. The clustering step based on the fraction of common contacts (FCC) [46] enhanced the scoring for TI and LI scenarios, achieving Top10 SR up to 65% in bound antigen runs and 55% in unbound (Fig. 3) for the LI case. Alternative scoring with the VoroIF-jury method [47, 48] was also performed, resulting in a minimal improvement in the LI scenario SR for Top1, but not enough to account for the increased computational costs (see Supplementary Fig. 3).

To measure the impact of the Flexible Refinement on the model quality, we calculated the improvement in the fraction of native contacts (*F*_*nat*_) on all acceptable models. For that, we computed the Δ*F*_*nat*_ by calculating the difference between each model *F*_*nat*_ measured after and before Flexible Refinement. The Δ*F*_*nat*_ median values for all information scenarios for both bound and unbound antigen runs ranged between 0.04 and 0.09, with distributions skewed toward positive values with maximum improvements around 0.4. This shows that Flexible Refinement does improve the quality of most models in every scenario (Fig. 4).

**Figure 4:**
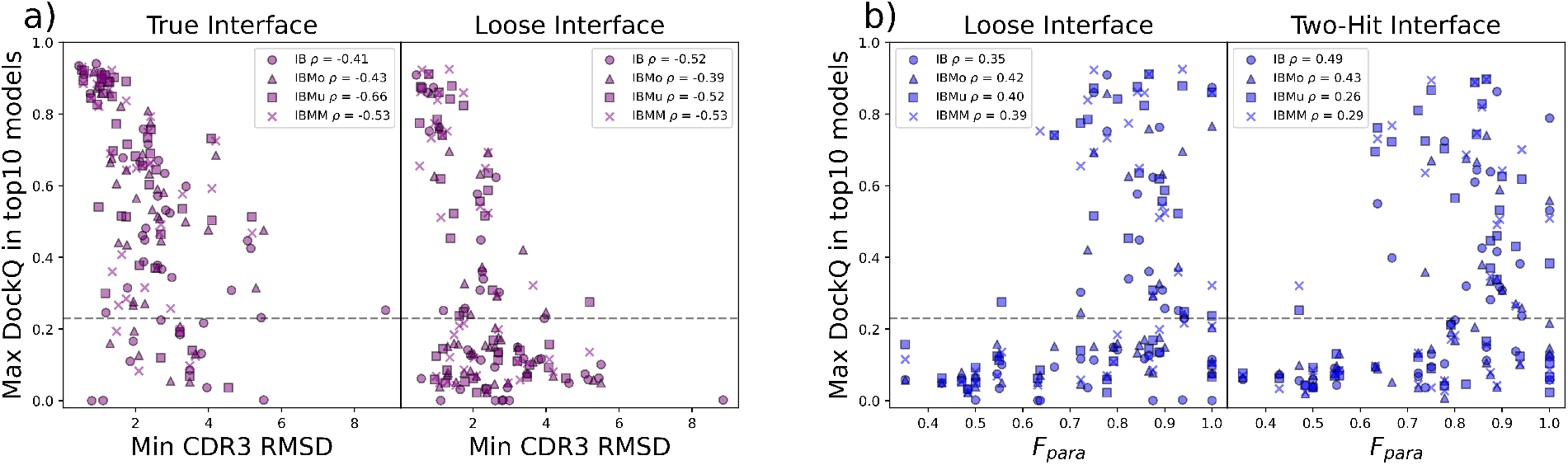
Nanobody structure quality and paratope definition are related to docking success. **a)** Scatter plot between the minimum CDR3 RMSD in the input models and the maximum DockQ achieved among the top10 models for each ensemble. Analysed models come from True Interface scenario and bound antigen. Pearson correlation coefficients for each ensemble are reported on the top right. The horizontal dashed line in each plot indicates a DockQ of 0.23, which is the cutoff used to define acceptable or better models. **b)** Scatter plots showing the relationship between the fraction of paratope present in the restraints (*F*_*para*_) and the maximum DockQ among the top 10 models for each ensemble. Data are shown for both Loose and Two-Hit Interface scenarios. Pearson correlation coefficients are plotted on the top left.

A detailed breakdown of HADDOCK results for all the nanobody-antigen complexes in the dataset is reported in the Supplementary Table 3.

### 2.5 CDR3 accuracy, CDR3 conformation, and paratope definition are critical features for docking success

We assessed the importance of the CDR3 loop accuracy on the quality of the docking results. The minimum CDR3 RMSD across the input nanobody ensemble strongly correlates with the maximum DockQ across the first ten models, reaching a maximum anti-correlation of -0.66 for the IBMu ensemble (Fig. 5a). We also observed a correlation between the maximum achieved DockQ and the epitope RMSD, which was a limiting factor for cases where the epitope RMSD is higher than 5Å (see Supplementary Fig. 4). These relationships between the accuracy of input models and docking success were previously described for antibody-antigen docking [34].

**Figure 5:**
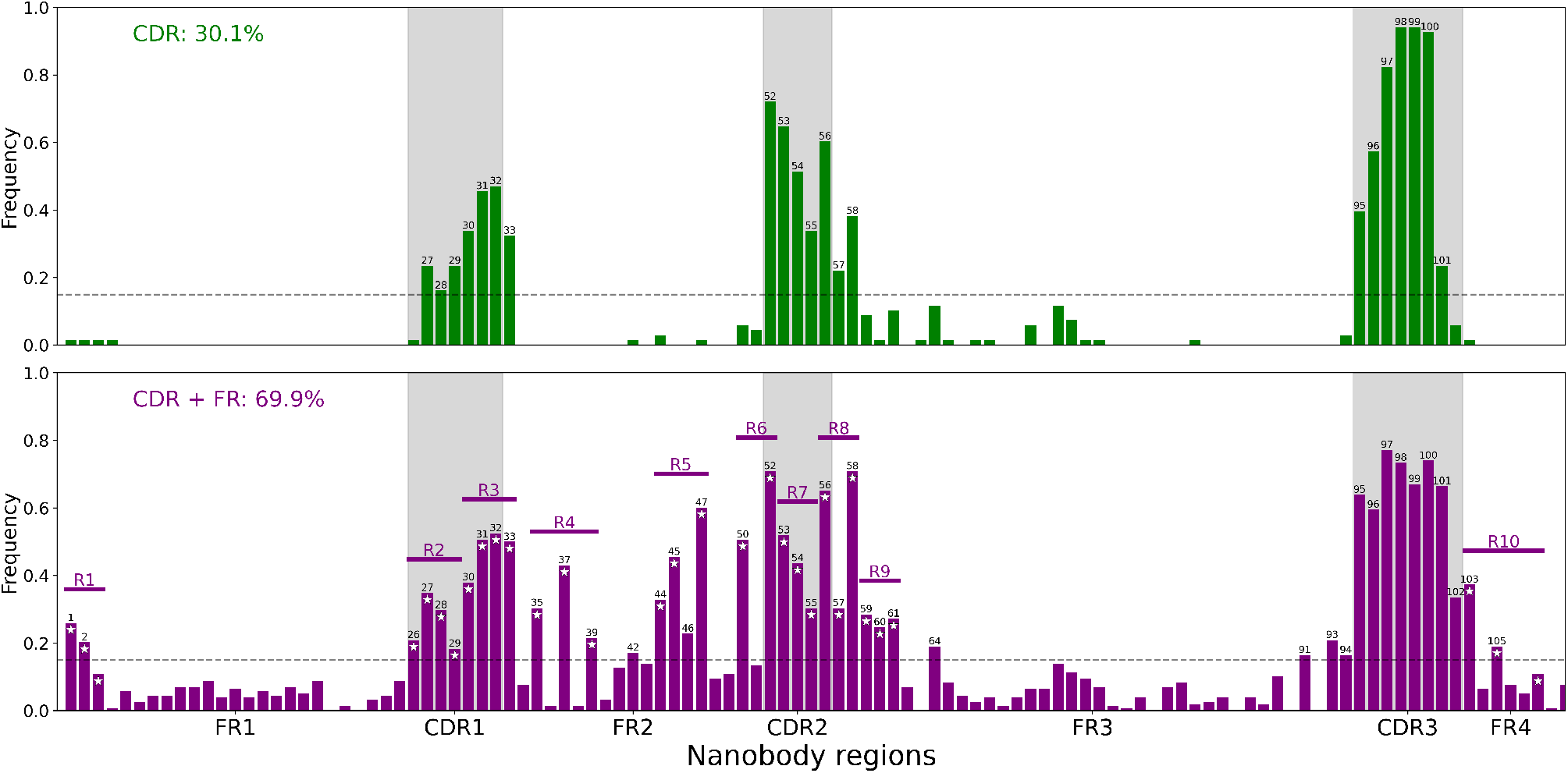
Paratope clusters. Bar plot showing the frequency of each residue in two cluster’s paratopes obtained after hierarchical clustering based on Jaccard distances. The top plot shows the frequencies for cluster 1 (CDR) and the lower plot for cluster 2 (CDR+FR). The CDR regions are highlighted in grey. Residue numbers with a frequency above 0.15 (dashed line) are plotted on top of the corresponding frequency bars. Only residue numbers based on the Chothia numbering scheme are considered for the frequency calculation and representation (e.g. residues 100A and 100B account as 100). Conserved regions (R1, R2, etc.) in the CDR+FR representation are shown with horizontal purple lines. White stars indicate the residues that are defined as paratope restraints inside of each of these regions.

The conformation (kinked or extended) of the CDR3 loop has a smaller but still relevant impact on the accuracy of the docking, as nanobodies with a kinked conformation consistently show better success rates, especially when the provided information is coarser (see Supplementary Fig. 5).

In the original HADDOCK recipe for LI, 2I, and AS information scenarios, the possible paratope residues of immune proteins are defined considering only the CDR regions. We found that some correlation exists between the fraction of the experimental paratope (*F*_*para*_) present in these restraints and the docking success (Fig. 5b): when less than 60% of the paratope is present in the docking restraints (i.e., located on the CDRs), HADDOCK struggles to generate a good model in the top 10 structures.

### 2.6 Framework-aware paratope description improves the modelling success

Motivated by the observed importance of *F*_*para*_ in the LI and 2I scenarios, we generated a new dataset of 226 nanobody-antigen experimental structures extracted from SabDaB following the same procedure used for the docking benchmark but without using any cutoff date (*Paratope Dataset*, see Methods).

We analysed the paratopes of these 226 complexes by using the classification criteria of ‘kinked’ and ‘extended’ CDR3 conformations based on dihedral angles as presented in Bahrami Dizicheh *et al*. [12]. When comparing the paratopes between these two classes, the only significant differences were found in the FR2 and FR3 regions, which were more frequently involved in the binding for the ‘extended’ conformations (see Supplementary Fig. 6). This is in discordance with the previously published results, which reported significant binding differences for CDR1, FR2 and FR4. Bahrami Dizicheh *et al*. [12] also reported a low frequency of FR3 involvement in antigen binding (*<* 35% of structures for both classes) and a higher frequency for FR1 (*>* 60%) on the binding, while we obtained less than 35% for FR1 and more than 70% for FR3.

With the aim of providing a more general analysis, we performed hierarchical clustering based on Jaccard distances over the *Paratope Dataset*. This allowed us to identify two main binding modes for nanobodies: the first resembles the one typical for antibodies and involves mainly CDR loops, explaining 30.1% of the dataset, while the second involves most frequently CDR3, in combination with regions of CDR1/2 and conserved framework regions, explaining 69.9% of the Paratope Dataset (Fig. 6). Based on these observations, we defined two possible paratope representations to be used as docking restraints: CDR (only the three CDR regions) and CDR+FR (CDR3 and some highly interacting CDR1, CDR2, and framework regions).

With the objective of better representing the nanobody-antigen interaction, we modified the LI and 2I restraints to divide equally the initially sampled structures into these two different paratope definitions (see Methods). We then ran the default pipeline with these new restraints using the best performing ensemble (IBMu) as a starting point.

For the LI scenario we obtained a maximum acceptable SR of 62.5% in bound antigen runs, and of 42.5% in unbound antigen runs for the Top10 models (see Supplementary Fig. 7). For the 2I scenario, the maximum SR achieved on Top10 models was 47.5% and 37.5% with bound and unbound Ag, respectively. The new paratope scenario managed to improve the LI scenario Top10 SR by 12.5% in bound antigen runs, while for the 2I scenario the SR were lower. This shows that expanding the nanobody paratope information can be beneficial for the docking performance, but not for all the approaches. Remarkably, when considering all models (Top200 SR), the LI scenario with modified paratope restraints and bound antigen structures achieves an acceptable SR of 95% (38 out of 40 successful structures). This illustrates how this paratope representation helps in generating acceptable models for almost the entire dataset.

## 3 Discussion

The prediction of nanobody-antigen complexes is of significant importance in the development of new therapeutical and diagnostic methods. This task is challenging for ML-based methods due to the large conformational space of CDR3 and the lack of coevolutionary signal in these interactions. In this work, we have built a non-redundant dataset of experimental nanobody-antigen structures, not seen by current AI prediction methods, with the aim of benchmarking state-of-the-art methods and identifying the best performing modelling protocol for this purpose.

The low success rate of AlphaFold2-Multimer on the dataset confirmed the limitation of this AI-based method for predicting nanobody-antigen complexes. The recently published AlphaFold3 [32] exhibits better performance, but it still struggles to identify the binding regions for several cases. This shows that information-driven docking algorithms remain a viable alternative when interface information is available and AI-based methods are failing.

We assessed nanobody structure prediction algorithms to determine the best possible input for docking. Methods generating multiple models such as ImmuneBuilder and AlphaFold2-Multimer generated the best models. The combination of predictions coming from different methods into ensembles further helped to improve the accuracy, increasing the likelihood to generate a good model for each case. We also assessed AlphaFold2-Multimer accuracy on antigen prediction, observing some limiting cases where this feature could doom docking to failure, especially for multiple chain antigens. In general, when homologous experimental structures of an antigen are available, they should be used instead of AlphaFold predictions.

To evaluate the influence of different levels of available interface information on the docking results, we ran HADDOCK with four distinct scenarios. The True Interface scenario condition allowed us to define the best achievable performance in terms of binding interface information (i.e. not considering here any specific distance restraints or contacts), reaching the highest SR (80% Top10) and highlighting how HADDOCK is able to generate correct models when detailed information is provided. The Loose Interface scenario might mimic the available information at the beginning of in silico design pipelines where some knowledge of the targeted epitope is available, while the Two-Hit Interface scenario simulates information coming, for example, from mutagenesis experiments. Both scenarios outperformed the AlphaFold2-Multimer baseline and performed similarly to AF3. This establishes the combination of information-driven docking and ML-modelled nanobody structural ensembles as a powerful approach to generate reasonable nanobody-antigen 3D models. Lastly, the All Surface scenario results showed markedly lower performance than other information scenarios, stressing the difficulty of predicting the correct epitope with docking [49]. When no information is available about the epitope, our recommendation is always to try AlphaFold and similar methods as the first option, only falling back to docking as a last resort.

To investigate the key determinants of docking success, we analysed the relationship between DockQ and different quality measures of the starting ensemble and docking data. We observed a strong correlation between the quality of the best available nanobody structure in the ensemble and the docking success (Fig. 5a). This is in line with the observed ability of HADDOCK to retrieve the best predictions among the ensemble and rank them higher based on the physics of the interaction. We highlight that using nanobody structures coming from AlphaFold3 in this pipeline is likely to increase the SR for all the presented scenarios, especially if a higher number of seeds is used at inference stage. The fraction of true paratope represented in the docking restraints, *F*_*para*_, also affects the modelling success. More specifically, docking with only the CDR loops as possible paratope amino acids was not representative of the true nanobody paratope for several cases within the dataset. With the objective of defining the paratope in a more realistic, framework-aware manner, we proposed two definitions of the nanobody paratope to drive the modelling. Using these two sets of restraints within the same docking run was observed to improve the results in the Loose Interface scenario. Especially when considering all final structures (top200), those new paratope restraints resulted in very high SR for unbound antigen (85%) and nearly a 100% SR for bound Ag, highlighting their ability to provide a robust description of the paratope for almost the entire dataset. However, while correct models are generated, identifying those remains difficult. This finding underlines the need for better scoring functions to identify the near-native models. Overall, these observations pave the way to boosting the pipeline’s performance with more advanced definitions of nanobody amino acids involved in the interaction, potentially derived from paratope prediction algorithms [50].

In summary, in the presented work we have defined a benchmark dataset of nanobody-antigen experimental structures and presented an assessment of state-of-the-art nanobody and nanobody-antigen structure prediction ML-based methods. We have demonstrated the potential of using these ML-based methods for information-driven docking, proposing pipelines for nanobody-antigen complex structure prediction. Finally, we have analysed the role of CDR and framework residues in antigen binding, which allowed us to propose new ambiguous restraint definitions to improve the docking performance.

## 4 Methods

### 4.1 Benchmark dataset assembly

We constructed the benchmark dataset with Chothia-numbered [51] bound nanobody structures downloaded from the SAbDab Nano database [52, 53] on November 15th, 2023. We only considered structures deposited after September 30th, 2021 to ensure that they were not part of the training set of any of the assessed ML-based prediction methods. We excluded NMR structures and only selected complexes with a resolution lower than 3.0Å, requiring that no CDR loop residues were missing. All regions from the nanobody sequence were defined following the Chothia numbering scheme from the references.

We ran CDR3 sequence homology checks, with a 60% identity threshold, on all the nanobody structures deposited before the training cutoff and inside the benchmark dataset, to remove redundancy in the loop region. Antigens shorter than 25 residues were excluded. PDB 7te8 was excluded from the dataset as the antigen was another nanobody. PDB 7nbb was excluded because the antigen is a polyubiquitin chain whose modelling is challenging and out of the scope of this paper. The resulting benchmark dataset consists of 40 nanobody-antigen complex structures (Supplementary Table 1) and is available from https://github.com/haddocking/nanobodies. PDB files handling was performed in Python using pdb-tools [54], and sequence retrieval and alignment using the MDAnalysis package [55].

### 4.2 Nanobody-antigen complex structure modelling and accuracy assessment

Nanobody-antigen structures were predicted using ColabFold v.1.5.3 using AlphaFold2-Multimer v.2.3 [23, 56] and with the AlphaFold3 server [32]. AlphaFold2-Multimer v.2.3. predictions were obtained from FASTA files containing both nanobody and antigen sequences following the ‘multimer’ pipeline. We ran the AlphaFold3 predictions for 5 different seeds in the web server to generate the same number of models as for AlphaFold2-Multimer. We ranked the resulting models by AlphaFold-rank, a weighted measure that uses ipTM and pTM scores. In AlphaFold3 disordered regions and atom clashes are also included in the final score [32].

### 4.3 Single structure modelling and accuracy assessment

Nanobody structures were predicted from FASTA files using AlphaFold2 v.2.3 (via ColabFold batch) [22, 56], ImmuneBuilder [27], RaptorX-Single [29] and NanoNet [28]. AlphaFold2 was used following the ‘monomer’ pipeline. ImmuneBuilder’s predictor NanoBodyBuilder2 default pipeline was modified to refine the top 4 models. AlphaFold2-Multimer v.2.3 nanobody predictions were extracted from the nanobody-antigen complex predictions.

We aligned the predicted and the reference structure on the framework residues prior to calculating RMSDs across different regions. RMSD calculations were performed for backbone atoms only, using the McLachlan algorithm [57] as implemented in Profit v.3.3 [58]. Antigen models were extracted from AlphaFold2-Multimer complex predictions in order to save computational time on antigen prediction. We aligned the predicted and the reference structure on all residues to calculate the global and epitope RMSD values.

### 4.4 Nanobody models clustering

We calculated the CDR3 RMSDs between all nanobody model pairs after fitting the structures on the framework residues. The resulting RMSD matrix was used for complete linkage hierarchical clustering using a 2.5Å threshold and a maximum number of clusters of 20 (never reached). Clustering was carried out with the scikit-learn Python package v.1.5.2.[59]. Within each cluster, the model with the best H3 loop confidence—measured by plDDT for AlphaFold or RMSPE for ImmuneBuilder (IB)—was selected as the cluster center. When models from IB and AlphaFold are in the same cluster, we always chose the AlphaFold model as the cluster center.

We performed a short energy minimization of models coming from AlphaFold to fix artefactual gaps that may appear in the generated structures using the HADDOCK3 [60] “emref” module (see example script in Supplementary Fig. 8). All predictions were formatted to a common numbering with pdb-tools.

### 4.5 Unbound antigen selection

We performed a sequence search in the PDB to find similar antigens alllowing for a maximum of one substitution, two missing residues or two insertions in order to include the antigens. We changed manually 7q6c-A PDB sequence homologue residue N84 to A84 to reverse the substitution. Antigens from 7tpr-C, 7wpf-C and 7whi-G were shortened due to their high length (*>* 400 residues) to a 200 residues selection including the epitope. For the remaining of the structures, we calculated the antigen global plDDT from the pool of AlphaFold2-Multimer complex predictions and selected the one with the highest value for each case. We then applied a short energy minimization only to the antigens coming from AlphaFold to fix artefactual structure gaps (see Supplementary Fig. 8). Visual inspection of PDB 7qne revealed that the relative orientation between chains A and E was incorrectly predicted by

AlphaFold2-Multimer, thus inherently limiting any downstream docking. Accordingly, we considered this case as failed in all unbound antigen docking runs.

### 4.6 Information scenarios and restraints generation

We assessed four different information scenarios to generate ambiguous restraints to drive the docking process using HADDOCK3 [60]:

- *True Interface*. We calculated the true interface of both the nanobody and antigen using a contact distance cutoff of 4.5Å in the experimentally determined structure. We set all residues as active for the docking: each active residue, when not present in the interface of a model, generates a restraint energy penalty [35]. No passive residues were used.
- *Loose Interface*. All the surface-exposed residues on the nanobody CDR loops are labelled as active. For the antigen, the epitope is defined combining the true epitope residues with the neighboring surface amino acids defined using a distance cutoff of 4Å (a residue is selected if any of its heavy atom is within the distance cutoff from any heavy atom of true epitope residue). This creates a wider region to be targeted by the nanobody. We treated all the selected antigen residues as passive in this case (contrarily to active residues, a passive residue not present in the interface will not generate an energetic penalty).
- *Two-hit Interface*. All the surface-exposed residues on the nanobody CDR loops are labelled as active. For the antigen, the only epitope information we use are two ‘hit’ residues, identified (e.g. by mutagenesis) to be key for the interaction. Those ‘hit’ residues were defined by running the “alascan” HADDOCK3 module on the experimental structures and selecting the two residues with the highest contribution to the interaction energy. The two ‘hit’ residues are set as active, with the neighbouring (within a cutoff distance of 7.5Å) surface-exposed residues are defined as passive.
- *All Surface*. All the surface-exposed residues on the nanobody CDR loops are labelled as active, and all antigen residues with a relative accessible surface area (RSA) ≥ 40% are defined as passive.

### 4.7 HADDOCK3 docking protocols

We used the previously published HADDOCK docking workflow for antibody-antigen complexes [33] for the first three scenarios. This workflow consists of the following steps:

1. Rigid body docking phase.
2. Selection of the top 200 models.
3. Flexible refinement phase.
4. Energy minimization.
5. Fraction of common contacts (FCC) clustering [46].
6. Selection of the top 4 models from each cluster.

The default sampling for step 1 is 1000 models (effective sampling of 10000 models, but only 1000 written to disk). For the All Surface scenario, we increased rigid body sampling to 10000, performed FCC-clustering and selected the top 10 models from each cluster for further flexible refinement instead of selecting the top 200 models (step 2, see Supplementary Fig. 9 for an example workflow). We ranked all models and clusters with the default HADDOCK scoring function, which, for the Flexible Refinement stage, is a weighted sum of various energies and buried surface area (BSA) calculated for each model complex:

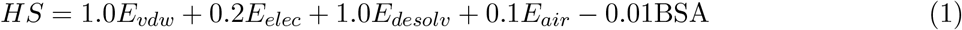

where *E*_*vdw*_ and *E*_*elec*_ are the intermolecular van der Waals and electrostatic energies, respectively, and *E*_*desolv*_ is an empirical desolvation energy term [61]. *E*_*air*_ is the restraint energy [35].

Alternative VoroIF-jury scoring [47, 48] was performed using a special branch of HADDOCK3 implementing the “voroscoring” module.

All the workflows were run using 12 cores on our local cluster (running on AMD EPYC 7451 processors). The average execution time (AET) of a HADDOCK3 workflow slightly depends on the scenario, with 2I runs (AET = 37 minutes) being faster than TI and LI scenarios (AET = 40 and 45 minutes, respectively). Runs with mixed restraints require roughly 30-40% additional computing time.

### 4.8 Model quality assessment

To assess the quality of the generated complex models we classify them following the CAPRI criteria [36, 37]. These criteria allow to assess the models’ similarity to the native structure by using the interface root mean square deviation from the reference (i-RMSD), ligand root mean square deviation (L-RMSD) and fraction of native contacts (*F*_*nat*_) parameters. Models are classified into

- high accuracy (*F*_*nat*_ ≥ 0.5 and either L-RMSD *<*= 1Å or i-RMSD ≤ 1Å)
- medium accuracy (*F*_*nat*_ ≥ 0.3 and either L-RMSD ≤ 5Å or i-RMSD ≤ 2Å)
- acceptable accuracy (*F*_*nat*_ ≥ 0.1 and either L-RMSD ≤ 10Å or i-RMSD ≤ 4Å)
- or incorrect

We define the acceptable success rate (SR) as the percentage of cases in the dataset with at least one acceptable or better quality predicted model in the TopN ranked models. We define the medium SR as the percentage of cases with at least one medium or better quality predicted model.

### 4.9 Paratope analysis and framework-aware restraints definition

The Paratope Dataset was assembled using the same criteria as the benchmark dataset, but without considering a training cutoff date. We excluded the structures belonging to our original benchmark dataset so as to avoid bias in our conclusions. The final dataset consists of 226 nanobody-antigen structures. For those, we calculated the paratope as the list of residues within a 4.5Å distance cutoff from the antigen (following the Chothia numbering scheme).

We performed the ‘kinked’ and ‘extended’ classification of the structures based on *α*101 and *τ* 101 dihedral angle calculations as described in Bahrami Dizicheh et al. [12]. We retrieved 135 ‘kinked’ structures, 60 ‘extended’ structures and 31 classified as ‘other’. Angle calculations were performed with the MDAnalysis package.

For clustering, we calculated a distance matrix with Jaccard distances computed between each paratope. Then we performed ward linkage clustering for a number of clusters N = 2 using the scipy Python package [62].

The new CDR+FR paratope was defined based on the observed conserved regions in the CDR+FR paratope cases. For these restraints, each residue in CDR3 was always defined as active, and several other regions were also defined as active (see below). When a region is defined as active, it is sufficient for any of the selected residues to be in contact with the antigen’s defined epitope for the restraint to be satisfied. The defined regions according to the Chothia numbering were:

- Region 1: residue 1, 2 or 3;
- Region 2: residue 26, 27, 28 or 29.
- Region 3: residue 30, 31, 32 or 33.
- Region 4: residue 35, 37 or 39.
- Region 5: residue 44, 45 or 47.
- Region 6: residue 50 or 52.
- Region 7: residue 53, 54 or 55.
- Region 8: residue 56, 57 or 58.
- Region 9: residue 59, 60 or 61.
- Region 10: residue 103, 105 or 108.

## 5 Data and Software Availability Statement

The input structures and restraint files used for the docking experiments and the numerical data and scripts used for generating the figures are available in the following Github repository: https://github.com/haddocking/nanobodies. HADDOCK3 is available from PyPI (https://pypi.org/project/haddock3) and GitHub (https://github.com/haddocking/haddock3). A tutorial on using HADDOCK3 for predicting nanobody-antigen complexes is available at https://www.bonvinlab.org/education/HADDOCK3/HADDOCK3-nanobody-antigen/

## 6 Acknowledgements

Financial support from Horizon Europe and from the European High Performance Computing Joint Undertaking, projects BioExcel (823830 and 101093290), and from the Netherlands e-Science Center (027.020.G13) is acknowledged.

