## Supplementary material for "Combining AI structure prediction and integrative modelling for nanobody-antigen complexes"

**Supplementary Table 1**. Benchmark dataset PDB IDs, CDR sequences and heavy chain species.

| PDB | Nb Chain | CDR1 | CDR2 | CDR3 | Heavy Chain Species |
|---|---|---|---|---|---|
| **7tgf** | B | GFTLDDY | SSSDGR | EEVCTLGIFGHGPDDY | *Vicugna pacos* |
| **8emz** | B | GRFFSSY | SWSGGS | AREGAYYPDSYYRTVRYDY | *Vicugna pacos* |
| **8h3x** | A | GFTFSRY | NSDGGR | PKRTYVPPSQFDD | *Vicugna pacos* |
| **8gz5** | B | GRTSSVY | TGNGGT | GGWGKERNYAY | *Vicugna pacos* |
| **8cwu** | B | GRTFSTS | DWSNTN | GGWGLTQPISVDY | *Lama glama* |
| **7x2m** | B | GDTLDLY | SPSGSR | SRPSAHYCSHYPTEYDD | *Vicugna pacos* |
| **7voa** | A | GFTLDYY | SGSGGI | VSHTVVAGCAFEAWTDFGS | *Vicugna pacos* |
| **7ubx** | C | GFTFSDY | NTDGS | GRVISASAIRGAV | *Camelidae* |
| **7th2** | C | PSLDYY | TGSEGS | ADPLPLVCTWGDEYDY | *Vicugna pacos* |
| **8en2** | D | GLTFSTN | NWNGGN | KMGRRLAVSRTLEEYDF | *Vicugna pacos* |
| **8ew6** | V | GFTFDDY | RIFDRH | GSFWACTRPEGAMDY | *Lama glama* |
| **7yz9** | B | GIILSIN | TAGGS | PAGRVGGT | *Vicugna pacos* |
| **7w1s** | B | ISFSSF | GIGG | WRMNDY | *Vicugna pacos* |
| **7zmv** | G | GSIFSIN | TSGGS | YGTYTLAPTGEGEYDDY | *Lama glama* |
| **7unz** | A | GSIFSTN | TSGSS | AGATIDLADFGS | *Vicugna pacos* |
| **8en3** | C | GRTDSES | NWRYAT | RYIYGSLSDSGSYDN | *Vicugna pacos* |
| **7tgi** | B | GLTLDYY | SSSDGR | DRDRLPSAITYEYNY | *Vicugna pacos* |
| **7nxx** | B | GGDTRPYITY | YTGGSG | GNGALPPGRRLSPQNMDT | *Camelus dromedarius* |
| **8dtn** | E | GFIFDSY | TSSGSS | LDYVIDGY | *Lama glama* |
| **8h3y** | E | GQTFSAW | NWNGER | MMGTYYSGSPKN | *Vicugna pacos* |
| **7q3q** | B | GLTFSSV | RWKFGN | ARVGEIIAVLISPSNYAY | *Vicugna pacos* |
| **7tpr** | E | GYTYSIC | NADGSN | HGTYDKYAPCGGFAGTYTY | *Camelus dromedarius* |
| **7x2j** | B | GRTFSRY | EWDGGT | GGNQYYSATYSIWNEYDF | *Vicugna pacos* |
| **7b5g** | D | GSIFSSN | SSRGDN | GSFYRGNYYGGSS | *Lama glama* |
| **7x2l** | B | GSISTLN | TLDGS | ENGGFFYY | *Vicugna pacos* |
| **8dtu** | A | GSTFSGY | TSSGAS | LDEGYLDYDS | *Lama glama* |
| **8dqu** | B | GLAFSMY | ISSGDS | PKFRYYFSTSPGDFDS | *Lama glama* |
| **7uia** | B | GFTFDDY | DSWSIN | EDRLGVPTINAHPSKYDYNY | *Vicugna pacos* |
| **7t5f** | F | GSIDSLY | QDGGS | KSTISTPLS | *Camelidae* |
| **7wki** | B | GLTVDDY | SSSNGS | AVSPNLECGTGPFGIYASYYGMDY | *Vicugna pacos* |
| **7x7e** | A | GGTLASF | DVINR | HFVPPGSRLRGCLVNELYNY | *Lama* |
| **7qbg** | E | GFTPGIY | SSRGSS | IYQPSNGCVLRPEYSY | *Vicugna pacos* |
| **7wd2** | D | GFTLDYY | SSNNS | EPDYSGVYYYTCGWTDFGS | *Vicugna pacos* |
| **7qne** | G | GRTFTAY | DQGRI | GAGFWGLRTASSYHY | *Lama glama* |
| **7r24** | E | GRTFSAY | SWSGNS | RKPMYRVDISKGQNYDY | *Vicugna pacos* |
| **7r24** | C | GRTSGAL | WWNDGT | RTPSSQTLY | *Vicugna pacos* |
| **7xqv** | B | GDTWWSS | SFYPTEY | IAWGPWMRTSWY | *Camelus bactrianus* |
| **7whi** | G | DFYFDYY | SGLGGA | RSPFGDYAFSY | *Homo sapiens* |
| **8bb7** | D | GFRLDYY | SSSGGS | SSYNTQRAECYGMDY | *Camelidae* |
| **8en0** | D | GRTFSSY | TGSGDS | YRTGGPPQ | *Vicugna pacos* |

**Supplementary Table 2.** Benchmark dataset antigen species and PDB from the used homologous antigens for docking. AF Multimer indicates the antigens that were modelled using AlphaFold2-Multimer.

| PDB | Ag Chain | Antigen Species | Antigen Name | PDB (chain) used for docking |
|---|---|---|---|---|
| **7tgf** | A | *Ricinus communis* | Ricin Chain A | 7y07 (A) |
| **8emz** | A | *Norovirus* | Gii.17 P Domain | 5lkk (A) |
| **8h3x** | C | *Bacteroides fragilis* | Fragilysin | AF Multimer |
| **8gz5** | A | Severe Acute Respiratory Syndrome Coronavirus2 | Spike Protein S1 | 7sxt (A) |
| **8cwu** | A | Severe Acute Respiratory Syndrome Coronavirus2 | Spike Protein S1 | 7z8o (A) |
| **7x2m** | E | Severe Acute Respiratory Syndrome Coronavirus2 | Spike Protein S1 | AF Multimer |
| **7voa** | B | Severe Acute Respiratory Syndrome Coronavirus2 | Spike Glycoprotein | 7eam (A) |
| **7ubx** | B | *Clostridioides difficile* | Toxin A | 7u1z (A) |
| **7th2** | A | *Ricinus communis* | Ricin A Chain | AF Multimer |
| **8en2** | A | *Norovirus* | Gii.10 P Domain | AF Multimer |
| **8ew6** | W | *Homo sapiens* | T-Cell Surface Glycoprotein Cd8 Alpha Chain | AF Multimer |
| **7yz9** | A | *Mycobacterium tuberculosis H37rv* | Adenylate Cyclase | AF Multimer |
| **7w1s** | A | Severe Acute Respiratory Syndrome Coronavirus2 | Spike Protein S1 | 7eam (A) |
| **7zmv** | A | *Homo sapiens* | Atp-Dependent Dna Helicase Q5 | AF Multimer |
| **7unz** | D | *Plasmodium falciparum* | Cysteine-Rich Small Secreted Protein Css, Putative | 7uny (A) |
| **8en3** | A | *Norovirus* | Capsid Protein Vp1 | 8emz (A) |
| **7tgi** | A | *Ricinus communis* | Ricin Chain A | 7y07 (A) |
| **7nxx** | A | *Homo sapiens* | Superoxide Dismutase [Cu-Zn] | 2c9v (A) |
| **8dtn** | F | *Homo sapiens* | B-Cell Lymphoma/Leukemia 11a | AF Multimer |
| **8h3y** | C | *Bacteroides fragilis* | Fragilysin | AF Multimer |
| **7q3q** | A | Severe Acute Respiratory Syndrome Coronavirus2 | Spike Glycoprotein | 7q3r (A) |
| **7tpr** | C | Severe Acute Respiratory Syndrome Coronavirus2 | Spike Glycoprotein | 7z8o (A) |
| **7x2j** | S | Severe Acute Respiratory Syndrome-Related Coronavirus | Spike Protein S1 | 7zh1 (A) |
| **7b5g** | C | *Escherichia coli K-12* | Lexa Repressor | 7ozj (A) |
| **7x2l** | E | Severe Acute Respiratory Syndrome Coronavirus2 | Spike Protein S1 | 7z8o (A) |
| **8dtu** | C | *Homo sapiens* \| *Lama glama* | B-Cell Lymphoma/Leukemia 11a \| Nanobody 5344n74d | 8dtn (F) |
| **8dqu** | F | Severe Acute Respiratory Syndrome Coronavirus2 | Non-Structural Protein 9 | AF Multimer |
| **7uia** | A | *Clostridium botulinum* | Neurotoxin Type E | 7ovw (A) |
| **7t5f** | D | *Clostridium botulinum* | Botulinum Neurotoxin Type B | 7na9 (A) |
| **7wki** | A | *Homo sapiens* | Complement Factor H | 3sw0 (X) |

| | | | | |
|---|---|---|---|---|
| **7x7e** | D | Severe Acute Respiratory Syndrome Coronavirus2 | Spike Protein S1 | 7z8o (A) |
| **7qbg** | C \| B | *Homo sapiens* \| *Homo sapiens* | Transcobalamin-2 \| Cd320 Antigen | AF Multimer |
| **7wd2** | B | Severe Acute Respiratory Syndrome Coronavirus2 | Spike Protein S1 | AF Multimer |
| **7qne** | A \| E | *Homo sapiens* \| *Homo sapiens* | Gaba(A) Receptor Subunit Alpha-1 \| Gamma-Aminobutyric Acid Receptor Subunit Beta-3 | AF Multimer (considered failed) |
| **7r24** | A | *Rattus norvegicus* | Activity-Regulated Cytoskeleton-Associated Protein | AF Multimer |
| **7xqv** | A | *Rattus norvegicus* | Rhoa | AF Multimer |
| **7whi** | A | Severe Acute Respiratory Syndrome Coronavirus2 | Spike Glycoprotein | 7x66 (R) |
| **8bb7** | B | *Mus musculus* | Plexin-B1 | AF Multimer |
| **8en0** | A | *Norovirus* | Capsid Protein Vp1 | 8emz (A) |

**Supplementary Table 3.** Alphafold modelling and docking results summary for the benchmark dataset. Alphafold and HADDOCK results are reported over the top 10 structures. HADDOCK results come from IBMu ensemble with unbound antigen, true interface scenario and Flexible Refinement stage.

| PDB | Nb Angle Class | Min CDR3 RMSD (Å) | Unbound Epitope RMSD (Å) | Max AF2 iPTM | Max AF3 iPTM | Max HADDOCK DockQ | HADDOCK Capri class |
|---|---|---|---|---|---|---|---|
| **7tgfB** | Extended | 1.738 | 0.463 | 0.88 | 0.90 | 0.819 | Medium |
| **8emzB** | Kinked | 2.192 | 1.256 | 0.15 | 0.11 | 0.669 | Medium |
| **8h3xA** | Other | 2.420 | 0.611 | 0.44 | 0.86 | 0.604 | Medium |
| **8gz5B** | Kinked | 1.020 | 0.441 | 0.27 | 0.85 | 0.814 | Medium |
| **8cwuB** | Kinked | 2.541 | 0.791 | 0.20 | 0.80 | 0.294 | Acceptable |
| **7x2mB** | Kinked | 1.470 | 1.124 | 0.36 | 0.18 | 0.245 | Acceptable |
| **7voaA** | Kinked | 1.740 | 0.557 | 0.38 | 0.14 | 0.400 | Acceptable |
| **7ubxC** | Extended | 0.989 | 4.607 | 0.20 | 0.91 | 0.516 | Acceptable |
| **7th2C** | Kinked | 2.693 | 0.393 | 0.54 | 0.16 | 0.362 | Acceptable |
| **8en2D** | Kinked | 1.171 | 0.398 | 0.750 | 0.820 | 0.587 | Acceptable |
| **8ew6V** | Kinked | 1.414 | 2.561 | 0.73 | 0.19 | 0.546 | Medium |
| **7yz9B** | Other | 3.285 | 1.191 | 0.16 | 0.86 | 0.182 | Incorrect |
| **7w1sB** | Failed | 1.365 | 0.681 | 0.40 | 0.25 | 0.723 | Medium |
| **7zmvG** | Extended | 1.066 | 1.680 | 0.23 | 0.70 | 0.566 | Medium |
| **7unzA** | Other | 4.543 | 1.036 | 0.17 | 0.23 | 0.127 | Incorrect |
| **8en3C** | Kinked | 2.684 | 1.753 | 0.17 | 0.11 | 0.311 | Acceptable |
| **7tgiB** | Kinked | 1.157 | 0.585 | 0.92 | 0.92 | 0.854 | High |
| **7nxxB** | Kinked | 2.562 | 0.565 | 0.19 | 0.18 | 0.391 | Acceptable |
| **8dtnE** | Other | 5.189 | 3.534 | 0.34 | 0.39 | 0.236 | Acceptable |
| **8h3yE** | Kinked | 0.565 | 3.575 | 0.60 | 0.15 | 0.635 | Medium |
| **7q3qB** | Kinked | 0.529 | 0.571 | 0.85 | 0.89 | 0.870 | High |
| **7tprE** | Extended | 2.374 | 2.192 | 0.25 | 0.36 | 0.331 | Medium |
| **7x2jB** | Kinked | 2.410 | 1.223 | 0.26 | 0.90 | 0.436 | Acceptable |
| **7b5gD** | Kinked | 1.240 | 0.964 | 0.21 | 0.16 | 0.766 | Medium |
| **7x2lB** | Extended | 0.784 | 0.779 | 0.86 | 0.79 | 0.721 | Medium |
| **8dtuA** | Kinked | 4.090 | 5.644 | 0.50 | 0.60 | 0.317 | Acceptable |
| **8dquB** | Kinked | 0.597 | 5.690 | 0.78 | 0.57 | 0.375 | Incorrect |
| **7uiaB** | Kinked | 3.485 | 2.220 | 0.21 | 0.60 | 0.079 | Incorrect |
| **7t5fF** | Extended | 1.778 | 0.570 | 0.26 | 0.11 | 0.641 | Medium |
| **7wkiB** | Kinked | 3.221 | 3.716 | 0.19 | 0.13 | 0.140 | Incorrect |
| **7x7eA** | Kinked | 1.283 | 0.875 | 0.66 | 0.18 | 0.586 | Medium |
| **7qbgE** | Kinked | 2.326 | 2.520 | 0.62 | 0.57 | 0.030 | Incorrect |
| **7wd2D** | Kinked | 1.617 | 0.579 | 0.25 | 0.85 | 0.283 | Acceptable |
| **7qneG** | Kinked | 2.106 | 13.376 | 0.62 | 0.80 | 0.130 | Incorrect |
| **7r24E** | Kinked | 1.113 | 1.589 | 0.86 | 0.87 | 0.478 | Medium |
| **7r24C** | Extended | 2.001 | 1.375 | 0.65 | 0.25 | 0.243 | Incorrect |

| | | | | | | | |
|---|---|---|---|---|---|---|---|
| **7xqvB** | Extended | 4.077 | 0.632 | 0.32 | 0.64 | 0.641 | Medium |
| **7whiG** | Kinked | 0.946 | 1.639 | 0.29 | 0.18 | 0.529 | Medium |
| **8bb7D** | Kinked | 1.387 | 1.206 | 0.18 | 0.13 | 0.399 | Acceptable |
| **8en0D** | Extended | 3.555 | 2.169 | 0.15 | 0.14 | 0.353 | Acceptable |

**Supplementary Figure 1. Violin plot showing the distribution of the backbone CDR3 RMSD from the best ranked nanobody prediction for the different methods**. The prediction methods shown in the plot are AlphaFold2-Multimer (AF Multimer), AlphaFold2 (AF Monomer), ImmuneBuilder (IB), RaptorX-Single (RXS), NanoNet (NN) and AlphaFold3 (AF3). Median values are indicated in the plot.

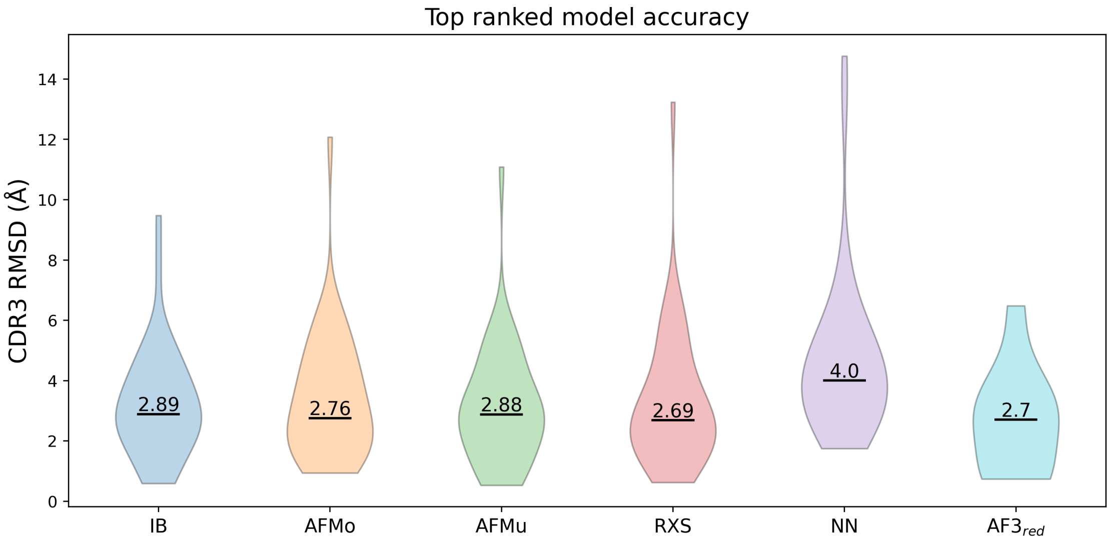

**Supplementary Figure 2. Bar plots showing docking success rate (SR) for All Surface scenario as a function of the TopN ranked structures**. The results shown are coming from the input ensemble of ImmuneBuilder + AlphaFold2-Multimer (IBMu) nanobody models and the bound antigen structures. The left plot shows results after Rigid Body stage of the pipeline, center left plot after Flexible Refinement (Flexref) stage and center right plot after Energy Minimization (Emref) stage. AlphaFold2-Multimer (AF2M) and Alphafold3 (AF3) results are shown in the right panel. The color scheme indicates the quality of the represented models following CAPRI criteria. From the plot we can observe how AF2M and AF3 outperform HADDOCK when no information is available and should be always preferred in this scenario. In terms of docking, the success rate is higher at the rigid-body docking stage.

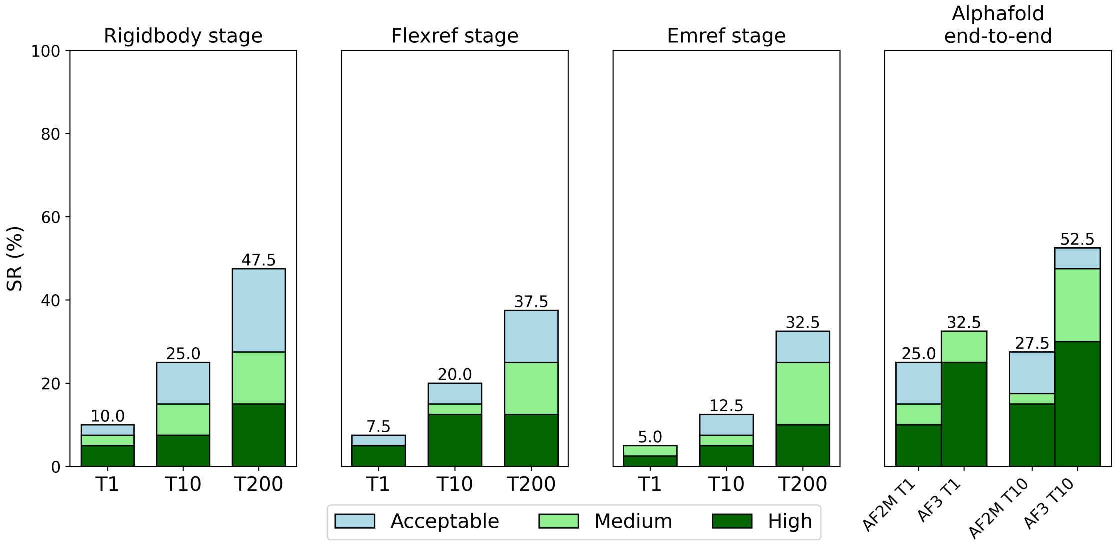

**Supplementary Figure 3. VoroIF-GNN scoring does not clearly improve the HADDOCK scoring of nanobody-antigen models**. Bar plot comparing model docking success rates (SRs) for TopN structures after Voronoi scoring (VS) and HADDOCK scoring (HS) for different information scenarios. Data shown comes from the input ensemble of ImmuneBuilder + AlphaFold2-Multimer nanobody models and unbound antigen structures. All considered models were scored after the Energy Minimization stage. The color scheme shows the quality of the represented models following CAPRI criteria.

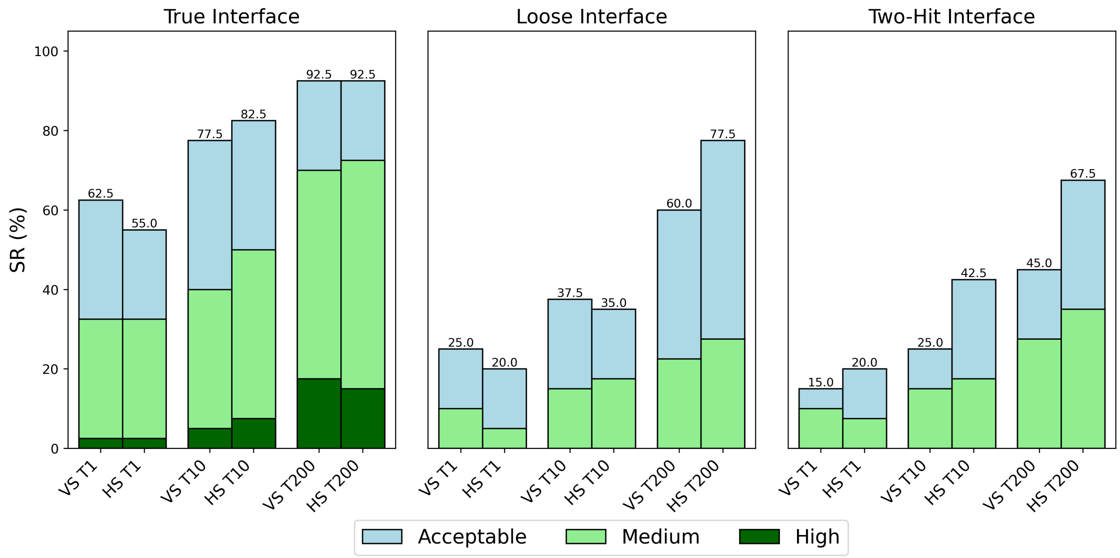

**Supplementary Figure 4. Scatter plot showing the relation between maximum DockQ among Top10 nanobody-antigen models and epitope RMSD**. The presented data come from the True Interface scenario runs with unbound antigen structures (excluding PDB ID 7QNE, for which unbound runs were not performed). The plotted models were retrieved after Energy Minimization stage. Represented nanobody structure ensembles are ImmuneBuilder (IB), ImmuneBuilder + AlphaFold2 (IBMo), ImmuneBuilder + AlphaFold2-Multimer (IBMu) and ImmuneBuilder + AlphaFold2 + AlphaFold2-Multimer (IBMM). Pearson correlation coefficients are shown for each ensemble. The horizontal dashed line indicates a DockQ score of 0.23 which is the cutoff for an acceptable model.

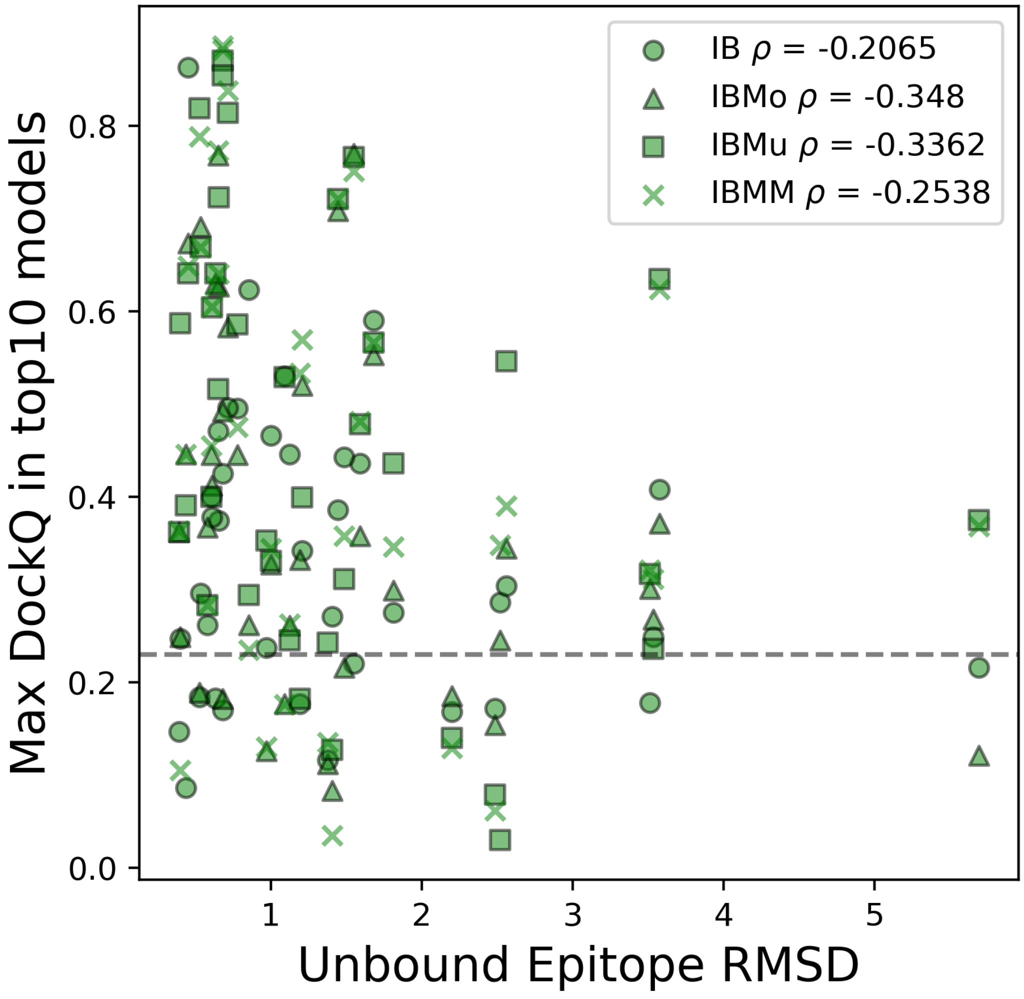

**Supplementary Figure 5. HADDOCK3 performance on nanobody-antigen docking for 'Kinked' and 'Extended' nanobodies.** From the original benchmark dataset 29 nanobodies are classified as 'Kinked' and 9 as 'Extended'. HADDOCK success rates (SRs) for nanobody-antigen docking models after Flexible Refinement stage for True, Loose, and Two-Hit Interface scenarios. Docking success is reported for the ensembles ImmuneBuilder (IB) and ImmuneBuilder + AlphaFold2-Multimer (IBMu), both for bound (B) and unbound antigens (U) for Top 10 ranked models. For IBMu we report also the cluster-based success rate. Clusters were defined based on Fraction of Common Contact (FCC) clustering of the models at the end of the workflow. Success is considered when at least one of the four first-ranked models in any of the Top10 ranked clusters is classified as acceptable/medium/high quality. The AlphaFold2-Multimer (AF2M) and AlphaFold3 (AF3) SRs for Top10 structures are provided on the right. Light blue, light green, and dark green indicate acceptable, medium, and high-quality models, respectively, as defined by the CAPRI criteria. Acceptable SRs (%) are shown on top of the bars.

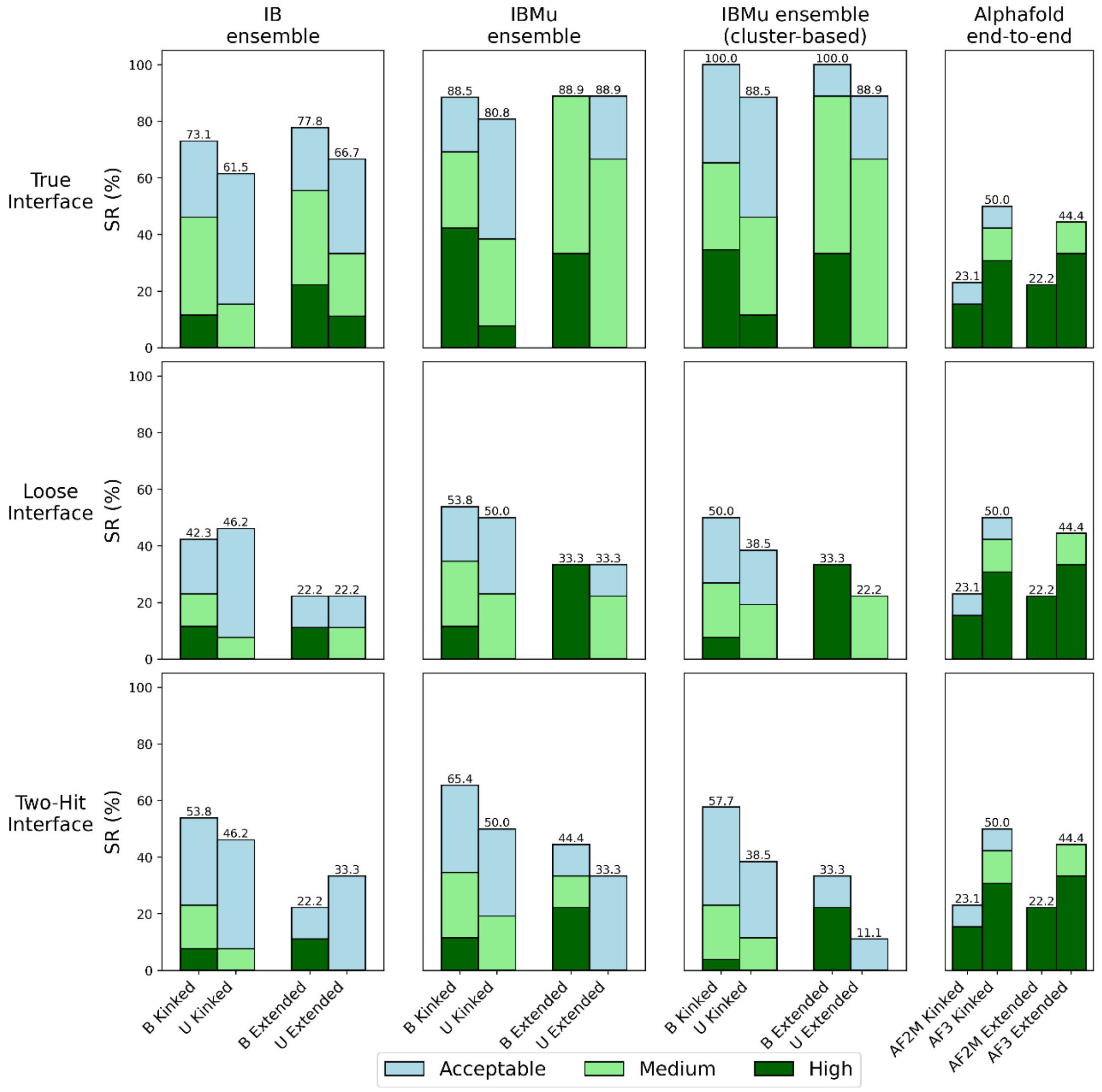

**Supplementary Figure 6.** Percentage of ‘kinked’ and ‘extended’ CDR3 conformation structures that involve each nanobody region in antigen binding, calculated over the Paratope Dataset (see main text). Structures that did not belong to any of the two classes are not considered in this analysis. Chi-squared significance in contingency tests between both classes for each region is labelled ($p$-values FR2: 0.0015; FR3: 0.012).

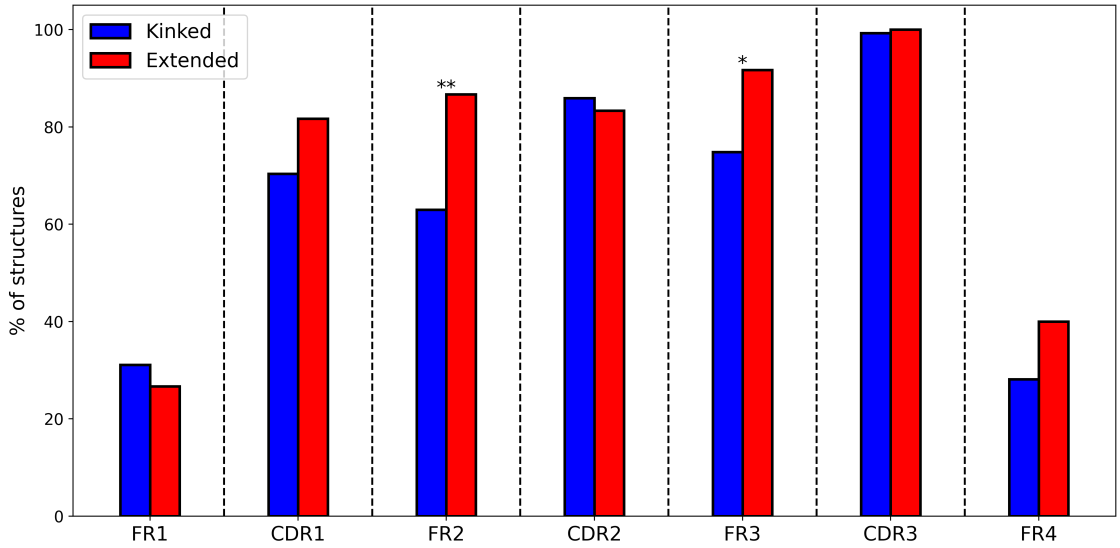

**Supplementary Figure 7.** Bar plots showing the docking success rates for the IBMu ensemble applying the mixed paratope restraints, namely combining CDR-only restraints (only the three CDR regions are defined as active) and CDR+FR restraints, where CDR3 and some highly interacting CDR1, CDR2, and FR regions are defined as active in HADDOCK. The two columns show Loose Interface and Two-Hit Interface scenarios. The color scheme shows the quality of the represented models following CAPRI criteria. Acceptable SR is plotted on top of the bars.

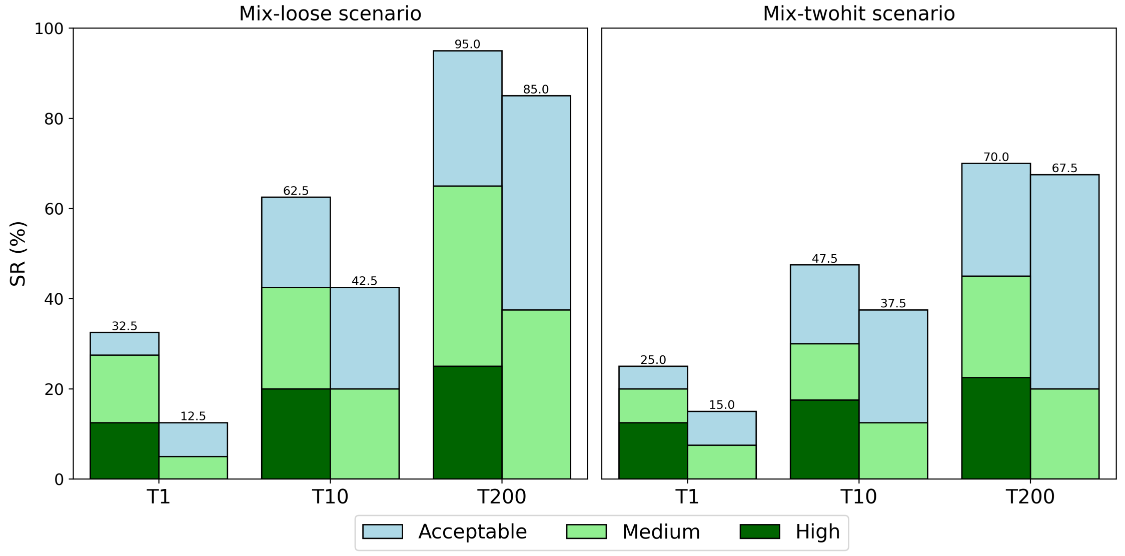

**Supplementary Figure 8.** Example workflow for the energy minimization refinement of nanobody or antigen structures from AlphaFold methods.

```
# ====================================================================
run_dir = “emref_7q3qB_af_multimer_centroids"
molecules = "7q3qB_af_multimer_centroids_ensemble.pdb”
clean = false
# ====================================================================
[topoaa]

[emref]
```

**Supplementary Figure 9.** Example workflow of the default nanobody-antigen HADDOCK pipeline.

```
# ==================================================================

run_dir = "nanobody_antigen_docking"
mode = “local”

molecules = ["7q3qB_ib_af_multimer_ensemble.pdb”,
             “7q3q_B_A_alphafold2_multimer_antigen.pdb"]
# ==================================================================
[topoaa]
[rigidbody]
tolerance = 5
ambig_fname = "7q3qB_real_ambig.tbl"
[caprieval]
reference_fname = " 7q3q-B:A_ref.pdb"
[seletop]
select = 200
[caprieval]
reference_fname = " 7q3q-B:A_ref.pdb"
[flexref]
tolerance = 5
ambig_fname = "7q3qB_real_ambig.tbl"
[caprieval]
reference_fname = " 7q3q-B:A_ref.pdb"
[emref]
tolerance = 5
ambig_fname = "7q3qB_real_ambig.tbl"
[caprieval]
reference_fname = " 7q3q-B:A_ref.pdb"
[clustfcc]
[seletopclusts]
top_models = 4
[caprieval]
reference_fname = " 7q3q-B:A_ref.pdb"
# ==================================================================
```

### Supplementary Method: Statistics and reproducibility

We calculated statistical differences using Chi-squared test using adjusted p-values. Significance levels were labelled as: n.s (non-significant), *($p <= 0.05$), **($p <= 0.01$), ***($p <= 0.001$). Statistical tests were performed using scipy Python package [1].
